# Reducing sialylation in melanoma increases classical pathway-mediated complement activation

**DOI:** 10.64898/2026.05.06.723302

**Authors:** Magali Coccimiglio, Georgia Clayton, Elisa C. Toffoli, Tanja D. de Gruijl, Richard B. Pouw, Fabrizio Chiodo, Yvette van Kooyk

**Author notes:** Corresponding authors: Yvette van Kooyk and Magali Coccimiglio.

## Abstract

Based on the success in pre-clinical models, methods that reduce sialylation in tumors have progressed to clinical trials, as this improves anti-tumor cellular responses. Immune responses against cancer can also be mediated by soluble, non-cellular mechanisms, such as the complement system. Dysregulation of the complement cascade and hypersialylation are hallmarks found across tumor types. Sialic acids are known to interact with complement proteins. However, the downstream pathways involved in the regulation of the complement cascade when reducing sialylation in tumors remain unclear. Here, using human melanoma cell lines and patient samples, we show that metabolic or enzymatic targeting of sialylation directly increases the activation of the complement, enhancing C3 opsonization of tumor cells and the formation of the membrane attack complex. This is mediated by the classical pathway of the complement system, in line with increased binding of immunoglobulins to tumor cells when sialylation is impaired. Our work positions the complement cascade as a relevant anti-tumor response playing a role when sialylation is targeted for cancer treatment.

**Graphical abstract:** 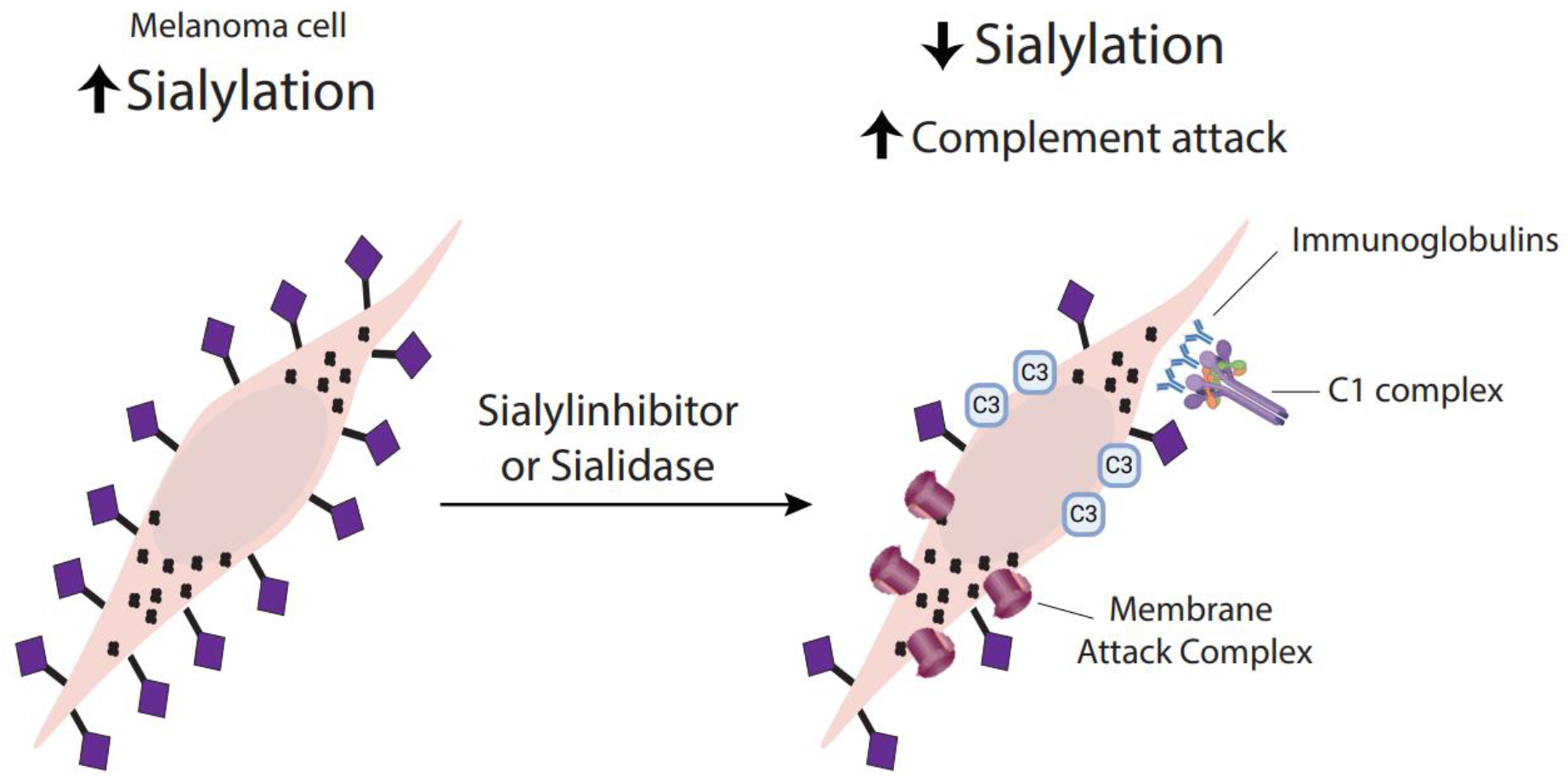

## Introduction

Tumors are very heterogeneous microenvironments, in which cancer cells interact with the immune system through a variety of complex mechanisms^1^. This interplay often leads to pro-tumor immunity, as tumor cells evade immune responses to be detected as “self”^2^. The increased expression of sialic acids, or hypersialylation, is a feature found across tumor types that leads to immune evasion and is related to worse patient survival and therapy response^3–5^. The sialylation pathway entails the addition of sialic acids to carbohydrate structures (glycans) on proteins, lipids, and RNA, catalysed by several enzymes called sialyltransferases^6,7^. These enzymes, localized in the Golgi apparatus, link sialic acids to other glycans in three different conformations, giving rise to a2,3-, a2,6-, and a2,8-sialylation. The sialylated glycoconjugates can then be transported to the cell membrane^6^. Sialic acids are usually added as the outermost glycans, making them an important structure for the interaction between cells and their environment. Different strategies that reduce sialylation increased anti-tumor immunity, favoured therapy response and delayed tumor growth in murine models^4,8^. This led to the first clinical trial (NCT05259696) aiming to reduce sialylation on tumor cells with an engineered fusion protein of a Fc portion with a sialidase (enzyme that cleaves off sialic acids).

Most work in targeting sialylation for cancer treatment, either using a metabolic inhibitor of this pathway or a sialidase fusion protein, was performed in mouse models and focused on the effects in cellular immunity^4,8^. Nevertheless, immune responses in cancer also include non-cellular mechanisms such as the complement system. Dysregulation of the complement cascades has emerged as a hallmark of cancer, impacting tumor progression, and anti-tumor cellular immune responses^9–11^. The complement system can be activated through three canonical pathways involving different molecules: the alternative, lectin and classical pathway^9^. Different cells and proteins in the tumor microenvironment mediate the activation of these pathways in tumors^12^. Previous research showed that sialic acids bind factor H (FH) and properdin, the main regulators of the alternative pathway^13–15^. Removal of sialic acids from cell surfaces was shown to increase C1 binding (the pattern recognition molecule initiating the classical pathway)^16^. However, these conclusions were mostly drawn using non-cellular approaches or non-carcinoma cells. In consequence, how targeting sialylation on human tumor cells affects the deposition of complement proteins and subsequent complement activation in cancer remains unclear.

Here, we used human melanoma cell lines and melanoma patient samples to study how targeting sialylation using different strategies affects complement activation. We used serum as a source for complement as well as complement blocking antibodies to modulate the different pathways of the complement system, dissecting which complement pathway is activated when sialylation on tumor cells is impaired.

## Results

### Tumor sialylation prevents C3 deposition and MAC formation

To determine the sialylation profile of three different human melanoma cell lines (MelJUSO, MelBRO, and MelAKR) by flow cytometry we used biotinylated recombinant proteins that bind either all surface sialic acids, or only a2,3-or a2,6-linked sialic acids. Additionally, an anti-poly-sialic acid antibody was used to detect a2,8-linked sialic acids. Melanoma cell lines expressed a2,3- and a2,6-sialic acids on their surface, while a2,8-sialic acids were undetectable (Fig. 1A). The expression of a2,3-sialic acids was higher than a2,6-sialic acids in all cell lines (Fig. 1B). After confirming that these cell lines are sialylated, we asked whether the presence of sialylation affects the activation of the complement cascade. Two strategies were used to reduce sialylation: a metabolic inhibitor (SI) and a neuraminidase (NEU), as previously described^17,18^. Cells non-treated (NT) or treated with SI or NEU were incubated with serum from healthy donors to determine C3 binding and the formation of the membrane attack complex (MAC, C5b-C9) on the tumor cells by flow cytometry as a measure of complement activation (Fig. 1C). We tested different concentrations of serum and found that 25% (v/v) gave the best difference in C3 deposition between serum incubation and the negative control (inactivated serum), without affecting cell viability (Supplementary Fig. S1A), hence, this concentration was used in further experiments. We first confirmed that SI and NEU treatments efficiently reduced surface sialylation on melanoma cell lines (Fig. 1D), without affecting the expression of complement regulators CD46, CD55, and CD59 (Supplementary Fig. S1B). We observed that reduction of sialylation, either with SI or NEU, significantly increased the deposition of C3 (Fig. 1E) and the formation of MAC (Fig. 1F) on tumor cells. To rule out the possibility of underestimating C3 deposition due to MAC-mediated cell death, we also performed the experiment adding eculizumab (a C5 inhibitor) to prevent MAC formation. Adding eculizumab efficiently decreased MAC formation without affecting C3 deposition or cell viability (Supplementary Fig. S1C-E) implying that our results on C3 binding are not affected by MAC deposition. Overall, our results demonstrate that reducing sialylation on tumor cells increases the activation of the complement cascade by promoting C3 deposition and MAC formation on tumor cells.

**Figure 1.**
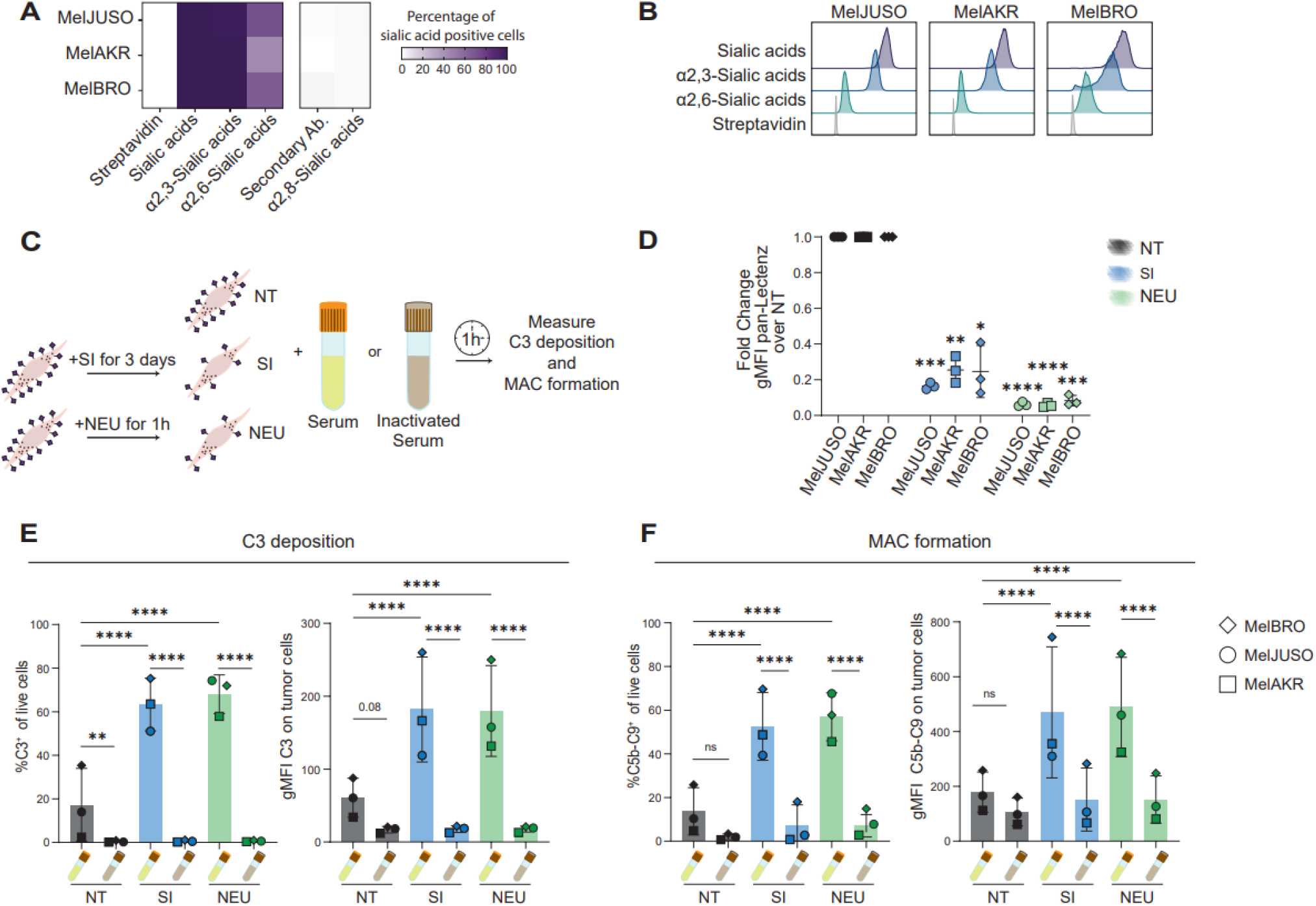
Tumor sialylation prevents C3 deposition and MAC formation. **(A)** Heatmap representing the percentage of sialic acid positive cells from live cells for three melanoma cell lines (MelJUSO, MelAKR, MelBRO) determined by flow cytometry. Data represents the mean of two independent experiments. Streptavidin and Secondary Antibody (Ab.) are the negative controls. **(B)** Histograms of sialic acids expression on the melanoma cell lines from (A). **(C)** Scheme of the complement activation experiment. Cells without treatment or after incubation three days with sialyltransferases inhibitor (SI) or one hour with neuraminidase (NEU), were incubated 1 hour with serum or inactivated serum. C3 deposition and membrane attack complex (MAC) formation were determined by flow cytometry. **(D)** Sialic acids expression on three melanoma cell lines after treatment with SI or NEU. Data shown as mean ± s.d. of the fold change geometric mean fluorescence intensity (gMFI) over the non-treated (NT) cells. Two-way ANOVA with Dunnett’s multiple comparisons test used for statistical analysis. **(E-F)** C3 deposition (E) and MAC formation (F) on NT, SI- or NEU-treated melanoma cell lines after incubation with 25% (v/v) serum or inactivated serum as percentage of positive cells from live cells (left) and gMFI C3 (right). Data shown as mean ± s.d. from seven independent experiments performed with two different serum pools. Nested one-way ANOVA with Tukey’s multiple comparisons test used for statistical analysis. ns = p_adj_>0.1, * = p_adj_ < 0.05, ** = p_adj_ < 0.01, *** = p_adj_ < 0.001, **** = p_adj_ < 0.0001.

### Reducing tumor cell sialylation promotes the activation of the classical pathway

To dissect which of the complement activation pathways drives complement deposition when sialylation is impaired in tumor cells, we performed the same experiment as in Fig. 1C with addition of anti-complement antibodies to modulate these pathways.

The alternative pathway is initiated by the spontaneous hydrolysis of C3, binding factor B (FB), which, after activation by factor D (FD), forms the C3 convertase C3bBb (Fig. 2A). This pathway is inhibited by factor H (FH) (Fig. 2A). The lectin pathway is activated by the binding of lectins, like mannose-binding lectin (MBL), complexed with MBL-associated serine proteases (MASPs), to carbohydrate structures (glycans) on cell surfaces (Fig. 2B). The classical pathway is initiated by the C1 complex (C1r, C1s and C1q), best known for its binding to and activation by the Fc portion of predominantly IgM and IgG, bound to their antigens (Fig. 2C).^9^

**Figure 2.**
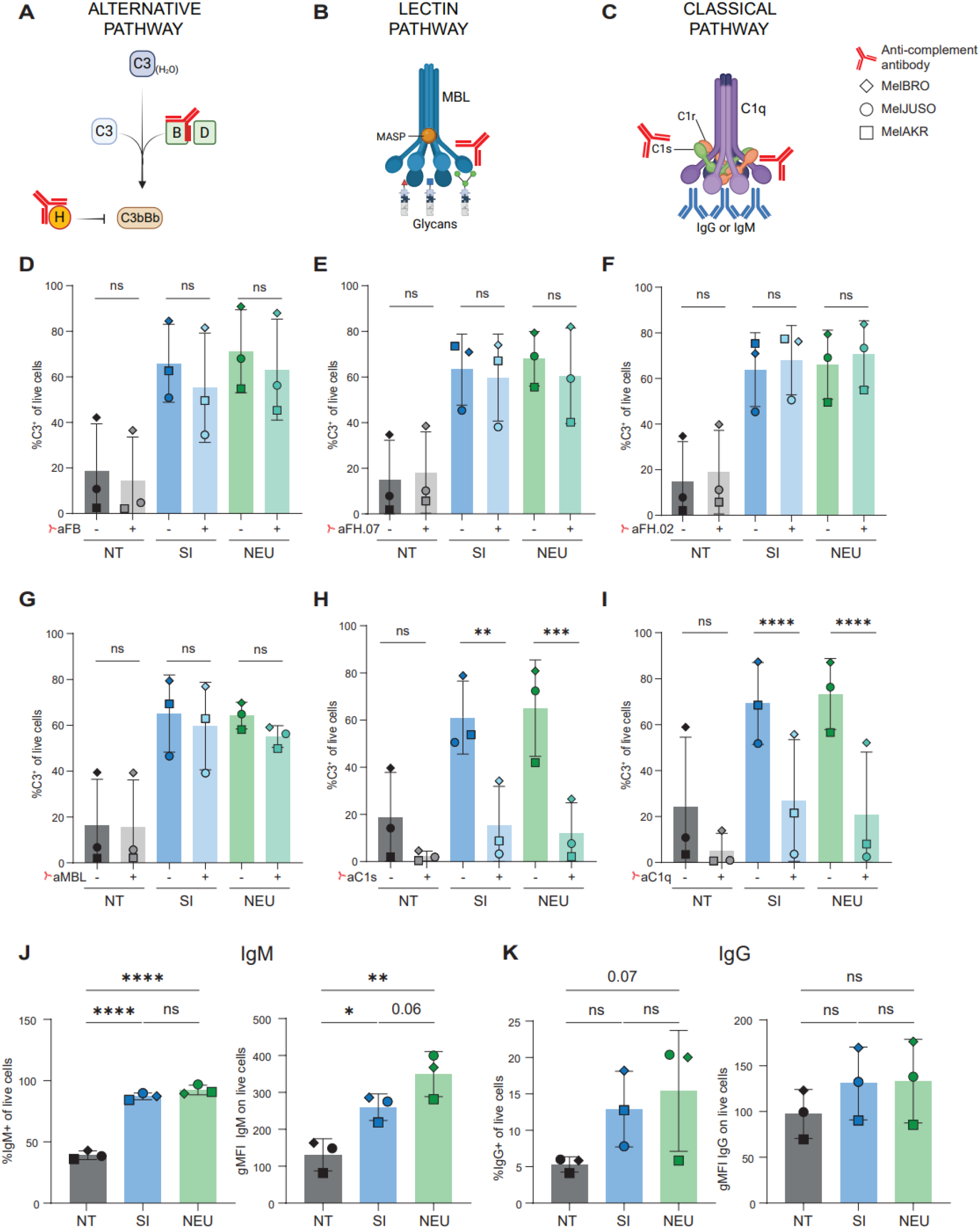
Reducing tumor cell sialylation promotes the activation of the classical pathway. **(A-C)** Representative diagrams of the three main pathways of activation of the complement system: alternative pathway (A), lectin pathway (B) and classical pathway (C). Red molecules indicate the anti-complement antibodies used to modulate the activation of each pathway. Images were modified from Biorender. **(D-I)** Binding of C3 to tumor cells after one hour incubation with serum, with or without anti-complement antibodies to modulate the alternative (D-F), lectin (G) or classical (H,I) pathways. Nested one-way ANOVA with Sidak’s multiple comparisons test used for statistical analysis. Antibodies used to modulate alternative pathway: aFB.01 (anti-factor B, blocks factor B activity), aFH.07 (anti-factor H.07, potentiates the activity of Factor H) and aFH.02 (anti-factor H.02, inhibits Factor H). Antibodies used to modulate lectin pathway: aMBL.01 (anti-mannose-binding lectin, blocks the mannose-binding lectin). Antibodies used to modulate classical pathway: aC1s (Sutimlimab, blocks C1s) and aC1q.89 (anti-C1q, blocks C1q). **(J-K)** Binding of IgM (J) or IgG (K) to tumor cells after 1 hour incubation with serum expressed as percentage IgM or IgG positive cells (left) or gMFI (right). Nested one-way ANOVA with Tukey’s multiple comparisons test for statistical analysis. gMFI = geometric mean fluorescence intensity. For all experiments cells were not treated (NT), treated with a sialylinhibitor (SI) or neuraminidase (NEU) prior to incubation with serum as described before. ns = p_adj_>0.1, * = p_adj_ < 0.05, ** = p_adj_ < 0.01, *** = p_adj_ < 0.001, **** = p_adj_ < 0.0001.

To modulate the alternative pathway, we used an antibody to block FB (aFB.01)^19^ and two antibodies to modulate the activity of FH: anti-FH.07 (aFH.07), which potentiates FH activity, causing inhibition of the alternative pathway, while anti-FH.02 (aFH.02) inhibits FH, leading to more alternative pathway activation^20^. We observed no change in C3 deposition with these treatments, either in NT, SI-, or NEU-treated tumor cells (Fig. 2D-F). The same results were observed when the lectin pathway was inhibited using anti-MBL.01^21^ (Fig. 2G). Notably, a strong reduction in C3 deposition on SI- and NEU-treated tumor cells was observed when the classical pathway was inhibited by blocking either C1s with sutimlimab, (Fig. 2H) or C1q with anti-C1q.89^22^ (Fig. 2I), with no effect in cell viability (Supplementary Fig. S1F,G). On NT cells, we observed the same trend, but it did not reach significance (Fig. 2H,I).

Activation of the classical pathway is mainly mediated by the binding of immunoglobulins (Igs) to cell surfaces, so we tested if reducing sialylation increases the binding of serum IgM and IgG to tumor cells. We found that IgM binding was significantly increased on SI- and NEU-treated melanoma cells compared to NT cells (Fig. 2J). As for IgG, we observed a non-significant trend to an increase in the percentage of IgG^+^ tumor cells when sialylation was reduced, with no changes in the density of IgG per cell (gMFI) (Fig. 2K).

Our data demonstrates that the classical pathway, and not the lectin and alternative pathways, drives the activation of the complement cascade on melanoma cells, especially when sialylation is impaired, likely attributed to increased binding of Igs.

### Targeting sialylation in melanoma patient samples promotes complement activation on tumor cells through the classical pathway

Our data demonstrates that when targeting sialylation on human melanoma cell lines, the classical pathway mediates the activation of the complement cascade on these cells. We then aimed to demonstrate if this also happens in a patient setting by utilising human melanoma samples. The first phase I/II clinical trial targeting sialylation in cancer was recently performed using a targeted sialidase, which aims to reduce tumor cell surface sialylation. Hence, we explored whether reduced sialylation of melanoma patient samples with sialidase (NEU), also increased the activation of the complement. Single cell suspension from melanoma patients, non-treated (NT) or treated with NEU, were incubated with serum to determine C3 binding and the formation of the MAC on tumor cells by spectral flow cytometry (Fig. 3A, Supplementary Fig. S2A,B). Reduction of sialylation by NEU (Supplementary Fig. S2C) increased C3 deposition (Fig. 3B) and MAC formation (Fig. 3C) on tumor cells. This was abolished when the classical pathway was inhibited using anti-C1q (Fig. 3D). In addition, 2/3 patients showed increased IgM binding to tumor cells (Fig. 3E), and all three patients had increased IgG binding to tumor cells after NEU treatment (Fig. 3F). Our data suggests that similar to the tumor cell lines, the classical pathway of the complement system is activated when sialylation in melanoma patient samples is reduced.

**Figure 3.**
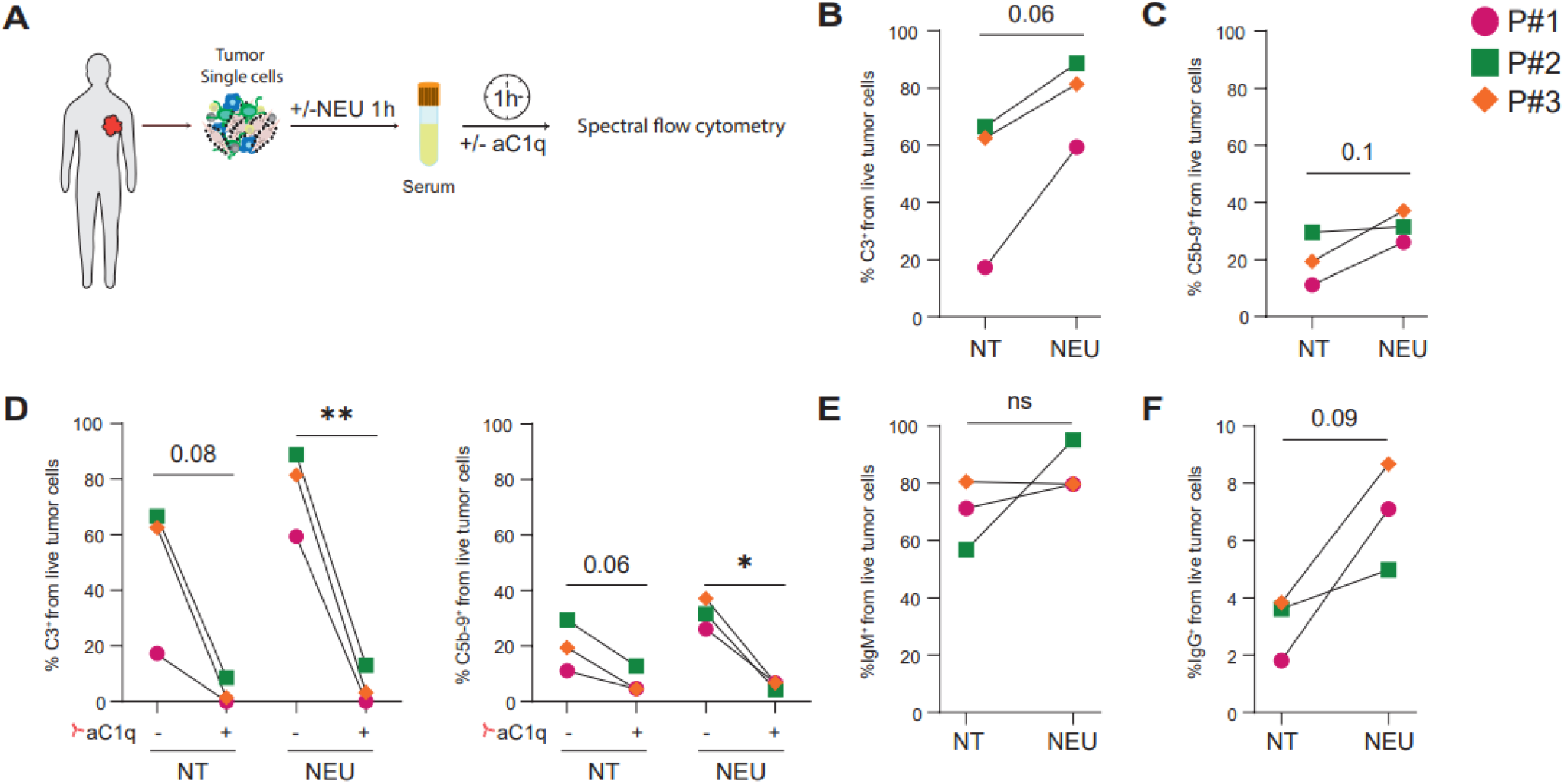
Targeting sialylation in melanoma patient samples promotes complement activation on tumor cells through the classical pathway. **(A)** Scheme of the experiment. Single cells from melanoma patient tumor biopsies (n=3) were treated with neuraminidase (NEU) for one hour and then serum was added at 25% (v/v) in the presence or absence of anti-C1q.89 (aC1q). After a one hour-incubation, cells were harvested and stained for spectral flow cytometry analysis. **(B-C)** C3 deposition (B) and MAC formation (C) on non-treated (NT) or NEU-treated melanoma cells from patient samples. Paired t-test used for statistical analysis. **(D)** C3 deposition (left) and MAC formation (right) on tumor cells from NT or NEU-treated melanoma patient samples, with or without the addition of aC1q. Paired t-test between aC1q + and – for each condition (NT and NEU). **(E-F)** IgM (E) and IgG (F) binding to tumor cells from melanoma patient samples. Paired t-test used for statistical analysis. P#= Patient number.

## Discussion

Here, we demonstrate that complement activation on melanoma cell lines and patient-derived tumor cells is enhanced when sialylation is reduced, promoting C3 binding and membrane attack complex (MAC) formation. This effect is driven by activation of the classical pathway, as its blockade abrogates the complement-activating consequences of decreased sialic acid expression. We show that immunoglobulins, mainly IgM, bind more strongly to tumor cells when sialylation is impaired, and may drive the activation of the classical pathway. Our work demonstrates that activation of the complement system is an important mediator of anti-tumor immunity when applying sialylation-targeting therapies.

Sialylation is emerging as a therapeutic target for many cancer types, including melanoma^3,6^. However, the majority of work has used animal models and focused on the effect in immune cells, while there is still limited data on patient material and the activation of innate soluble immune responses^4,18,23–25^. Although previous research has indicated that sialic acids regulate the activation of the alternative pathway by interfering with factor H and properdin binding^13–15,26,27^, our findings suggest it is largely the classical pathway of the complement system that is activated against melanoma cells and mediates the enhanced C3 binding when tumor sialylation is decreased. This is in line with previous reports showing that mainly genes of the classical pathway, rather than alternative or lectin pathways, are enriched in patient samples from several tumors, including melanoma^12^. Previous work also showed that when the classical pathway is efficiently activated, amplification via the alternative pathway is not required, supporting our results^19^. In addition, removing sialic acids from microglia favours C1q binding, which can increase classical pathway activation^16^. It is worth mentioning that we did not see differences in cell killing either blocking MAC formation or the classical pathway. This indicates that melanoma cells are resistant to MAC-mediated killing, and then C3 opsonization and consequent phagocytosis may be more relevant in the context of the tumor microenvironment to achieve cell killing^9^. This needs to be considered for current and future anti-cancer therapies targeting either C3 and/or MAC formation^9,12^.

Our work shows that immunoglobulins, mainly IgM, have increased binding to tumor cells when sialylation is impaired, on both cell lines and melanoma patient samples. As the classical pathway is strongly activated by IgM^28^, this can explain the enhanced complement activation through the classical pathway when sialylation is impaired. Future research is needed to elucidate the epitope specificity of the IgM antibodies, which are likely natural antibodies against glycan epitopes that may be better exposed when sialic acids are removed, enhancing their binding^29,30^. However, a role for IgG which was also found to be increased when sialylation is impaired could not be ruled out. One limitation of our study is that we used serum from healthy donors due to the lack of access to serum from melanoma patients. Previous studies have shown that anti-glycan antibodies are present in cancer patients and differ from healthy individuals, and that these immunoglobulins can increase complement-mediated cytotoxicity towards tumor cells^30–35^. Future research into the anti-glycan Igs repertoire in cancer patients and its correlation with glycan structures on tumor cells will shed light on the mechanisms behind glycan-dependent classical pathway activation. Our work opens avenues to investigate whether using immunoglobulins against glycans enhances complement activation and has a synergistic effect with sialylation-targeting strategies for cancer treatment.

Sialylated molecules are known to protect cells against complement activation, although this was mostly described in non-cancerous cells and looking only at cell lysis^15,16,36,37^. In the present work, we used a sialidase to target sialylation in melanoma samples from patients. This enhanced binding of C3 and MAC formation on tumor cells, which was mediated by the classical pathway, recapitulating our results with melanoma cell lines. Higher expression of classical pathway genes in melanoma correlates with better patient overall survival^12^. On the contrary, higher sialylation is linked to worse patient prognosis^4,5^. Our data shows that reducing sialylation increases the activation of the classical pathway in patient samples. Further research is needed to determine if this has a synergistic effect and further improve patient outcome.

Overall, our work offers a basis to not only focus on cellular immunity, but also consider the activation of the complement system as an anti-tumor mechanism playing a heightened role when sialylation is targeted in cancer.

## Methods

### Cell lines

The human melanoma cell lines MelBRO, MelAKR and MelJUSO were obtained from the Cancer Center Amsterdam. They were cultured in Iscove’s Modified Dulbecco’s Medium (IMDM, Gibco) supplemented with 10% Fetal Calf Serum (Biowest) and 1% penicillin/streptomycin (Gibco), at 37°C and 5%CO_2_with humidified atmosphere.

### Patient samples

Frozen single cell samples from melanoma patients were selected from a previous study of Active Specific Immunotherapy (ASI) with autologous whole-cell tumor vaccines, carried out at the VU Medical Center. Tumor dissociation and cryopreservation methodologies for these samples were previously reported^35^.

### Serum

All sera were obtained with informed consent according to the local ethics committees in accordance with Dutch law. Blood was obtained from healthy donors at Sanquin Research and the Amsterdam UMC by venipuncture according to local protocols in serum tubes. Clotting was allowed at room temperature (~22°C) for 30-60 min. Samples were centrifuged at 4°C, 1800x*g* for 10 minutes. After centrifugation, serum was separated and transferred to tubes on ice. Pools of serum were obtained mixing equal part of serum from different donors and aliquoted for storage at −80 °C until use.

The serum pools were incubated in a water bath at 56°C for 1 hour for inactivation of the complement system just before the experiments were performed.

### Anti-complement antibodies

Anti-C3.19, anti-CD46.01, anti-CD55.01, anti-factor B.01, anti-factor H.02, anti-factor H.07, anti-C1q.89 and anti-MBL.01 were previously developed and produced by Sanquin Research^19–22,38,39^. Anti-C5 was produced recombinantly by Evitria AG (Zurich, Switzerland) based on the sequence of Eculizumab (Alexion). Sutimlimab (anti-C1s, Sanofi) was obtained by collecting surplus from used Enjaymo™ injection bottles.

### Sialylation reduction treatments

For the cell lines, cells were harvested and plated in T25 flasks (400.000 cells/flask in 4 mL medium) and Sialyltransferase Inhibitor 3Fax-Peracetyl Neu5Ac (Sigma Aldrich, dissolved in DMSO at 200 mM stock) was added at 200 µM. Then they were incubated for 3 days at 37°C and 5%CO_2_ with humidified atmosphere.

For cell lines and patient single cells, cells were washed two times with Phosphate Buffer Saline (PBS) and incubated with neuraminidase (Sigma Aldrich, 1:250 in PBS) for 1h at 37°C and 5%CO_2_ with humidified atmosphere. To wash out the remaining neuraminidase, cells were washed twice with cell culture medium previous to their use for experiments.

### Sialic acids flow cytometry staining

Cells were washed with PBS, followed by staining with eBioscience Fixable Viability Dye eFluor 450 (Invitrogen) for 10 minutes. After washing with Hanks’ Balanced Salt Solution (Gibco) + 0.5% (w/v) Bovine Serum Albumin (BSA, Sigma Aldrich) (HBSS-BSA), cells were incubated with a mix of SiaFind Pan-Specific, Alpha 2,3-Specific or Alpha 2,6-Specific biotinylated Lectenz (2 µg/mL, LectenzBio) and Streptavidin-APC (Becton Dickinson) in HBSS-BSA; or with a mix of Anti-Polysialic acid [735], Rabbit IgG, Kappa (Absolute Antibody) and secondary donkey anti-rabbit IgG AlexaFluor647 (Invitrogen) in PBS + 0.5% BSA (PBA), for 30 minutes. Cells were washed once with PBS and fixated with 2% Paraformaldehyde in PBS (PFA) in PBS for 15 minutes. Cells were washed and resuspended in PBA for acquisition in the BD LSRFortessa X-20 cytometer. All incubations were performed at 4°C in the dark. FlowJo v10.10 was used for data analysis.

### Complement regulators flow cytometry staining

Cells were harvested and washed once with PBS for staining of dead cells with eBioscience Fixable Viability Dye eFluor 450 (Invitrogen) for 10 minutes. After one wash with PBA, cells were stained with a mix of anti-CD55.01-AF647, anti-CD46.03-AF555 or CD59 Recombinant Rabbit Monoclonal Antibody (108), PE (Invitrogen) for 30 minutes. After a wash with PBA and one wash with PBS, cells were fixed with 2% PFA for 10 minutes. All incubation were done on ice in the dark. Cells were washed with PBA and acquired in the BD LSRFortessa X-20. FACS Diva software was used for acquisition and FlowJo v10.10 for analysis.

### Complement activation experiments on cell lines

Cells were plated in 96-wells flat bottom cell culture plates (100.000 cells/well) in medium. Then serum or inactivated serum was added at the correct % (v/v) (indicated in figures). After 1 hour incubation at 37°C and 5%CO_2_, 10 µM EDTA was added to stop complement activity. For the modulation of the complement pathway, anti-complement antibodies were added at 150 µg/mL just before adding the serum.

C3 deposition and membrane attack complex (MAC) formation were determined using conventional flow cytometry. Cells were harvested and washed once with PBS for staining of dead cells with eBioscience Fixable Viability Dye eFluor 450 (Invitrogen) for 10 minutes. After one wash with PBA, cells were stained with a mix of anti-C3.19 AF488 (Sanquin Research, in house labeled) and Anti-C5b-9 + C5b-8 antibody [aE11] (Abcam) AF647 (in house labelled) for 30 minutes. After a wash with PBA and one wash with PBS, cells were fixed with 2% PFA for 10 minutes. All incubation were done on ice in the dark. Cells were washed with PBA and acquired in the BD LSRFortessa X-20. FACS Diva software was used for acquisition and FlowJo v10.10 for analysis.

### Immunoglobulins binding on cell lines

Cells were plated in 96-wells flat bottom cell culture plates (100.000 cells/well) in medium. Then serum or inactivated serum was added at the corresponding % v/v. After 1 hour incubation at 37°C and 5%CO_2_, 10 µM EDTA was added to stop complement activity.

Cells were harvested and washed once with PBS for staining of dead cells with eBioscience Fixable Viability Dye eFluor 450 (Invitrogen) for 10 minutes. After one wash with PBA, cells were stained with a mix of FITC anti-human IgM Antibody (Biolegend) and APC anti-human IgG Fc Antibody (Biolegend) for 30 minutes. After a wash with PBA and one wash with PBS, cells were fixed with 2% PFA for 10 minutes. All incubation were done on ice in the dark. Cells were washed with PBA and acquired in the BD LSRFortessa X-20. FACS Diva software was used for acquisition and FlowJo v10.10 for analysis.

### Complement activation experiments on patient samples

Cells were plated in 24-wells flat bottom plates (500.000 cells/well) in medium. Then serum or inactivated serum was added at 25% (v/v). After 1 hour incubation at 37°C and 5%CO_2_, 10 µM EDTA was added to stop complement activity. For the blocking of the classical complement pathway, anti-C1q.89 was added at 150 µg/mL just before adding the serum.

C3 deposition and MAC formation were determined using spectral flow cytometry. Cells were harvested and washed once with PBS for staining of dead cells with Fixable Viability Stain 780 (Becton Dickinson) for 10 minutes. After one wash with PBA, cells were incubated for 30 minutes with a mix of antibodies (Table 1) combined with and BD Pharmingen Human BD Fc Block (Becton Dickinson) and True-Stain Monocyte Blocker (Biolegend). All incubations were on ice in the dark. After washing with PBA, samples were acquired in the 5 Lasers Cytek Aurora. SpectroFlo software was used for acquisition, unmixing and compensation. FlowJo v10.10 was used for the rest of the analysis.

**Table 1.** Antibodies used for staining of patient samples for spectral flow cytometry.

| Antibody | Fluorochrome | Company |
| --- | --- | --- |
| Mouse anti-human CD45 | PE-Cy7 | Becton Dickinson |
| CD31 Antibody, anti-human, REAfinity | PE-Vio615 | Miltenyi |
| Human NG2/MCSP | PE | R&D systems |
| Anti-C3.19 | AF488 | In house, Sanquin Research |
| Anti-C5b-9 + C5b-8 antibody [aE11] | AF647 | Abcam (in house labeled) |
| anti-human IgM Antibody | BV785 | Biolegend |
| anti-human IgG Fc Antibody | Spark PLUS UV395 | Biolegend |

### Statistical analysis

GraphPad Prism software was used for statistical analysis. The tests used are described in the figure legends. P-values less than 0.05 were considered statistically significant. P values and number of replicates (n) are indicated in the figures.

