## Supplementary Figures for "Reducing sialylation in melanoma increases classical pathway-mediated complement activation"

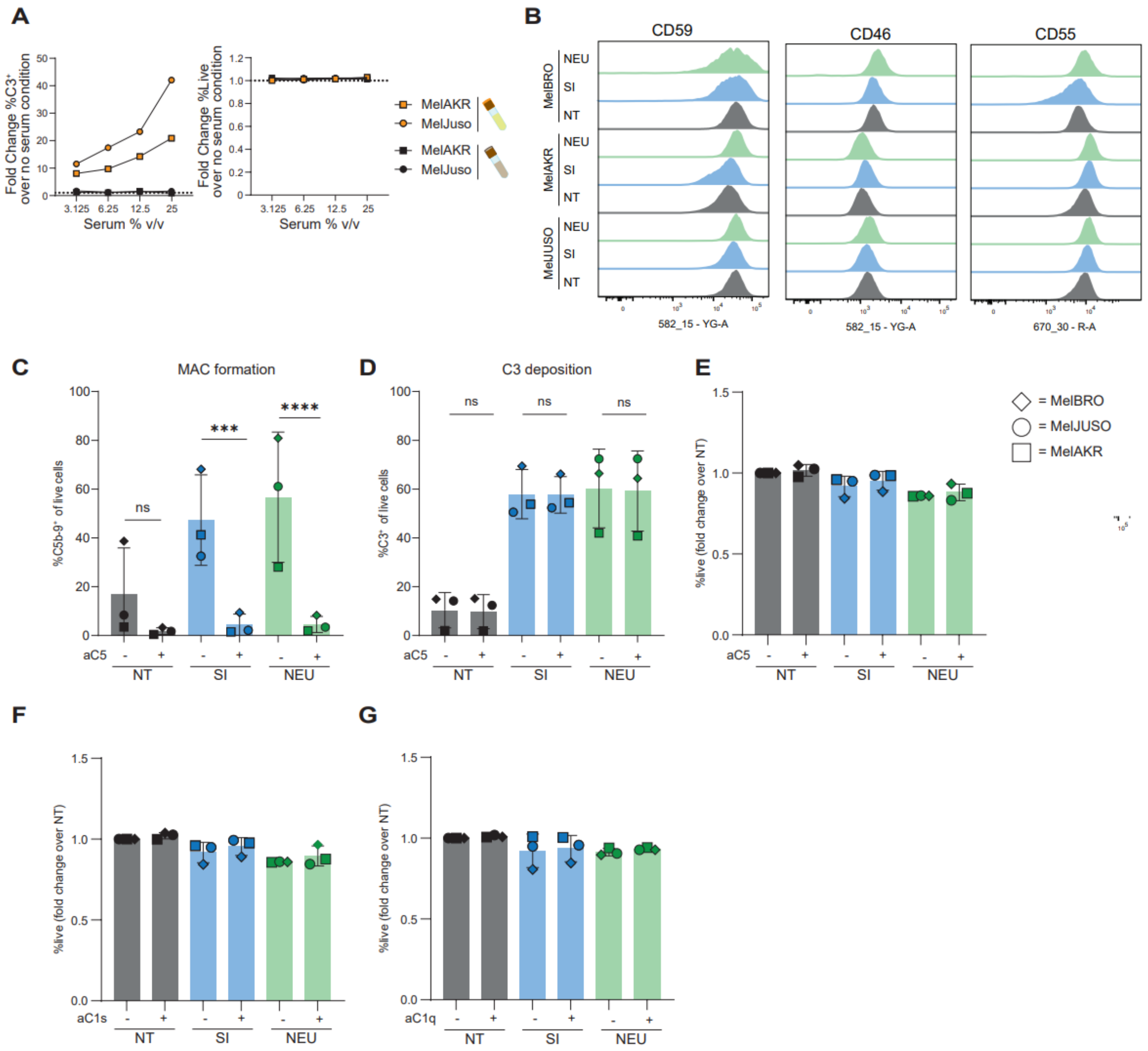


**Supplementary Figure S1. Tumor sialylation prevents C3 deposition and MAC formation. (A)** C3 deposition (left) and viability (right) of melanoma cells after 1 hour incubation with increasing amounts of serum or inactivated serum as described in Fig.1C. Data shown as fold change over the no serum condition. **(B)** Histograms showing the expression of complement regulators CD59, CD46 and CD55 on melanoma cell lines non-treated (NT) or after treatment with sialyltransferases inhibitor (SI) or neuraminidase (NEU). **(C,D)** MAC formation (C) and C3 deposition (D) on NT, SI- or NEU-treated melanoma cells after 1 hour incubation with 25% (v/v) serum in the presence or absence of eculizumab (aC5). Data shown as mean ± s.d. from three independent experiments. **(E-G)** Percentage of live cells from single cells on NT, SI- or NEU-treated melanoma cells after 1 hour incubation with 25% (v/v) serum in the presence or absence of eculizumab (aC5) (E), aC1s (F) or aC1q (G). Data shown as mean ± s.d. from three independent experiments. Nested one-way ANOVA with Tukey's multiple comparisons test used for statistical analysis. ns = $p_{adj}>0.1$, * = $p_{adj} < 0.05$, ** = $p_{adj} < 0.01$, *** = $p_{adj} < 0.001$, **** = $p_{adj} < 0.0001$.

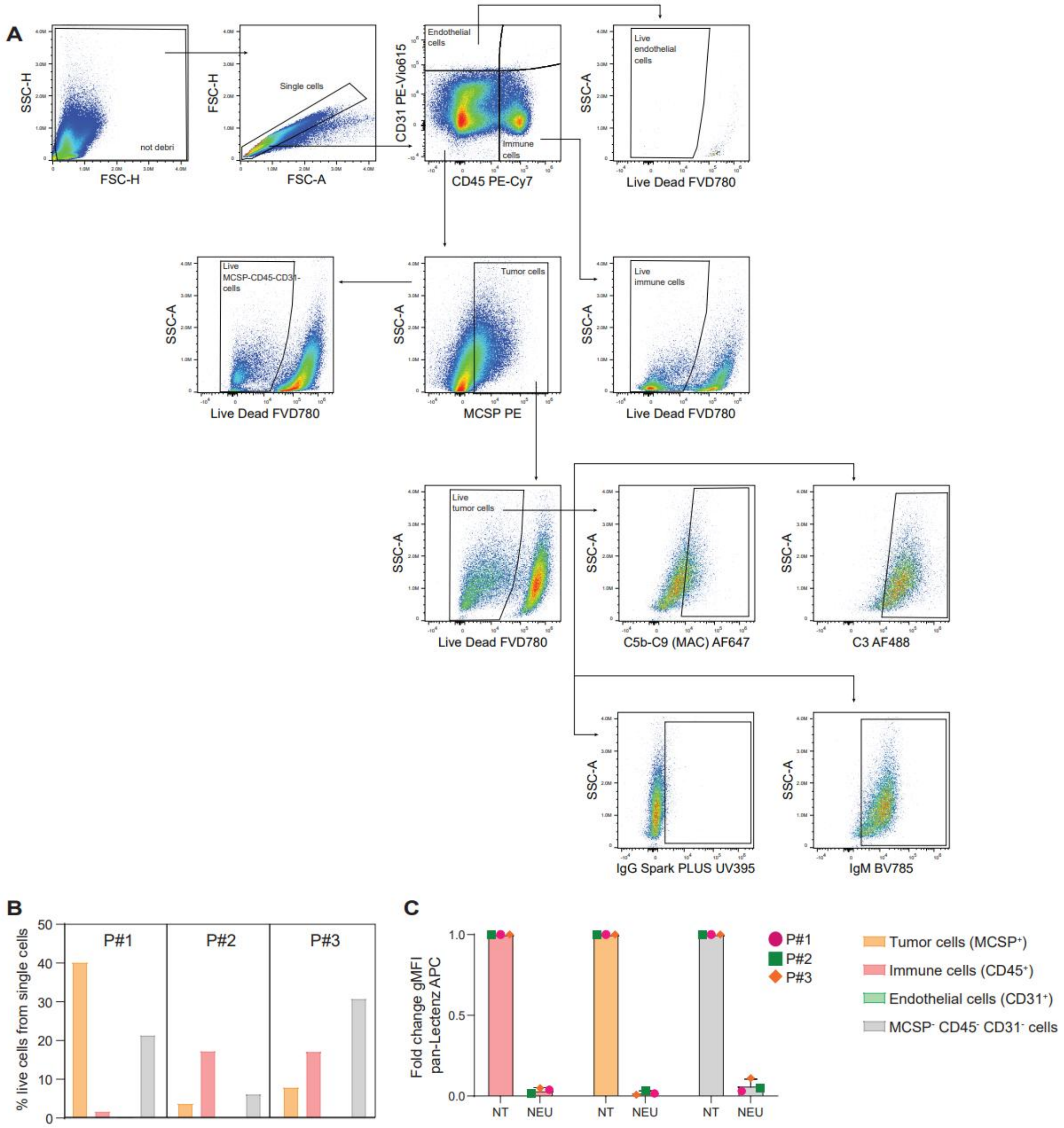


**Supplementary Figure S2. Targeting sialylation in melanoma patient samples promotes complement activation on tumor cells through the classical pathway. (A)** Gating strategy used for spectral flow cytometry panel on patient samples. The gates were set according to Fluorescence Minus One (FMO) controls and the no serum condition. **(B)** Frequency of tumor cells, immune cells, endothelial cells and the rest of cells in each patient sample. **(C)** Sialic acids expression on cells form patient samples after treatment with NEU. Data shown as fold change geometric mean fluorescence intensity (gMFI) over the NT condition. P#= patient number.
